# Where to refine spatial data to improve accuracy in crop disease modelling: an analytical approach with examples for cassava

**DOI:** 10.1101/2025.03.07.642020

**Authors:** Yevhen F. Suprunenko, Christopher A. Gilligan

## Abstract

Epidemiological modelling plays an important role in global food security by informing strategies for the control and management of invasion and spread of crop diseases. However, the underlying data on spatial locations of host crops that are susceptible to a pathogen are often incomplete and inaccurate, thus reducing the accuracy of model predictions. Obtaining and refining data sets that fully represent a host landscape across territories can be a major challenge when predicting disease outbreaks. Therefore, it would be an advantage to prioritise areas in which data refinement efforts should be directed to improve the accuracy of epidemic prediction. In this paper, we present an analytical method to identify areas where potential errors in mapped host data would have the largest impact on modelled pathogen invasion and short-term spread. The method is based on an analytical approximation for the rate at which susceptible host crops become infected at the start of an epidemic. We show how implementing spatial prioritisation for data refinement in a cassava-growing region in sub-Saharan Africa could be an effective means for improving accuracy when modelling the dispersal and spread of the crop pathogen cassava brown streak virus (CBSV).

## 1. Introduction

A major challenge when modelling epidemics of crop disease is the lack of complete and accurate maps of the distribution of susceptible crops across a landscape (1). The challenge is especially relevant when addressing epidemic threats to food production in climate-vulnerable regions. For example, in Ethiopia, the absence of high quality crop type maps is a challenge for modelling of the unfolding wheat rust epidemic (2,3) (but see the recently provided Ethiopian Crop Type 2020 dataset (4) for a single 2020/21 cropping season). In sub-Saharan Africa the modelling of the expanding epidemic of cassava brown streak disease (5) may be sensitive to local spatial uncertainty in crop distribution (6) and therefore the quality of model predictions relies on the quality of the available data on the spatial distribution of cassava (7). The problem of using incomplete or inaccurate crop maps is that they can amplify uncertainties in modelling the extent and rate of disease spread by either under- or over-estimating the crop area as well as failing to account for the spatial structure of crops (e.g. clustering of crops often at multiple scales with concomitant crop-free gaps). In cases when it is impossible to rectify incomplete and inaccurate data, models and methods that use partial data have been proposed (for example, in modelling the spread of livestock diseases when exact spatial locations and clustering of livestock holdings are unknown, see (8–12)). However, additional data collection is sometimes possible (e.g. by ground-based methods or via remote sensing (13–15)) and this could be used to refine incomplete mapped data with the purpose of improving the accuracy of epidemic modelling. The challenge then becomes how to use additional resources for data refinement most efficiently.

In this paper, we address the problem of finding geographic regions where new data on host crop distribution would lead to the strongest improvement of the accuracy in modelling the spread of crop diseases. We aim to develop a rapid analytical assessment method to estimate the impact of potential inaccuracies in crop data sets on the resulting predictions of epidemic spread derived from computer simulations of an individual-based model (IBM) for epidemic spread. Analytical approximations have been used to address the problems of incomplete host maps in studies of human and livestock by mapping local changes in the basic reproduction number *R*_0_ to assess regions at-risk (16), to design and implement vaccination strategies (17). Sellman *et al*. (12) also used local estimates of *R*_0_ to assess the effects of clustered disaggregation of county-scale livestock premises on epidemic predictions. Suprunenko *et al*. (18) developed an alternative approach based upon the infection rate, *r*, to predict the impact of the spatial structure of a crop landscape on epidemic dynamics of crop disease. The method involved calculation of the infection rate, *r* (18), which has the advantage of encompassing SI epidemics for which *R*_0_ is undefined, while also applying to SIR epidemics where *R*_0_ = *r*/*μ*, and *μ* is the removal or recovery rate of infected fields.

We now use the approach of Suprunenko *et al*. (18) to develop a method of spatial prioritisation of areas for new data. Our focus is on crop disease but the approach has wider applicability. For example, whereas the total number of livestock premises were known in the analyses of Sellman *et al*. (12) our method allows for uncertainty in the numbers of susceptible crops (analogous to livestock premises) across a landscape through which a pathogen is spreading.

We illustrate the application of the method to the analysis of the spread of cassava brown streak virus (CBSV) in an arbitrarily selected cassava production region in sub-Saharan Africa. Cassava is one of the most important staple food crops. An estimated 800 million people in Africa rely on cassava for their primary calorific intake (19). Cassava production in sub-Saharan Africa has come under increasing pressure due to the rapidly expanding range of CBSV (20–23). Recent work has provided improved maps of cassava production in sub-Saharan Africa (7) that are then used in combination with a parameterised epidemic model to predict spread and arrival times of CBSV throughout the region (5). We show how the method of our paper can be used to identify local areas within the host crop (cassava) landscape where potential errors in maps of cassava production would impact most strongly on predictions of CBSV spread. The analyses show that surveying and refining data in certain areas substantially improve accuracy in predictions of the epidemic spread model. Hence, correcting for insufficient or inaccurate datasets could make a crucially important contribution towards a better preparedness of regions to the on-going expansion of the CBSV pandemic.

## 2. Methods

### 2.1. Cassava data

The map with currently the highest spatial resolution for cassava production in sub-Saharan Africa is known as CassavaMap derived by Szyniszewska (7). We used the cassava production layer from CassavaMap (7) and converted it to discrete fields of cassava as detailed in Godding *et al*. (5) (i.e. one raster 1 km by 1 km cell can have a maximum of 1000 identical cassava fields) to create the data that describe the spatial distribution of cassava fields in sub-Saharan Africa. A 336 km by 336 km area was arbitrarily selected on the border between Cameroon and Central African Republic, the data describing this area were denoted as a host landscape H (Figshare (24), Fig. 1).

**Figure 1.**
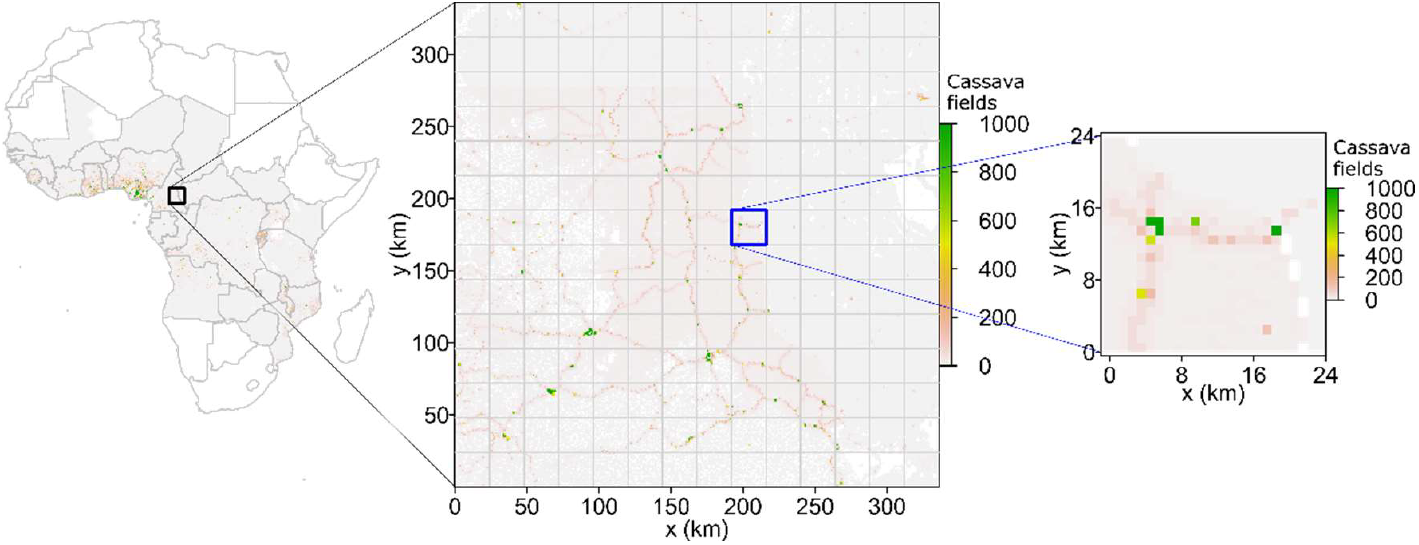
Host landscape. Data (as raster with spatial resolution 1 km by 1 km) on spatial distribution of cassava production extracted from CassavaMap (7) and converted to fields (5), see Methods; here we show the entire 336 km by 336 km landscape H (available online (24)). We resolved the 336 km by 336 km landscape into a square lattice with mesh size of 24 km. For convenience, the 24 km by 24 km area outlined by the blue boundary is used in Fig. 2 to illustrate the analyses that were conducted on the entire 336 km by 336 km landscape.

### 2.2. Potential inaccuracies in cassava data

Here, we briefly summarize likely inaccuracies in CassavaMap that stem from inaccuracies in the underlying data, as reported by Szyniszewska (7). The map was derived using administrative unit level data on cassava production and harvested area from 32 countries (7) and disaggregating the data according to a rasterised human population density model, LandScan 2014 (25). CassavaMap is available at a resolution of approximately 1 km by 1 km. According to Szyniszewska (7), errors in the spatial distribution in LandScan 2014 or in production statistics would result in errors in CassavaMap (7). Moreover, the size of human population in the current year may be very different from the size 2014 (25) thus affecting the estimate of the area of cassava fields. Lastly, due to coarse administrative granularity of input data on cassava production (e.g. the data from some countries were available only on a national scale (7)), some areas in CassavaMap have cassava production distributed in a flatter manner than in reality. Recent work by Hassall *et al*. (6) has raised some additional challenges in mapping cassava at landscape and regional scales. Here in the absence of further improvements or detailed mapping of extensive regions we treat CassavaMap as the best available starting point to illustrate the application of our method below.

For the purpose of this paper, we examine two possible scenarios related to the accuracy of crop maps. Firstly, where the actual spatial distribution of cassava fields is more clustered than presented in CassavaMap. To illustrate inaccuracies of this type we used a previously established method (18) of aggregating hosts locally: we considered the cassava landscape on a square lattice with mesh size of 2 km and assumed that cassava fields can be aggregated within each 2 km by 2 km cell.

The second scenario represents inaccuracy where the distribution of cassava fields was under-estimated in CassavaMap due to changing human population over time. We compared the human population distribution model, LandScan 2014 (25), used in CassavaMap and its most recent version, LandScan 2022 (26). The comparison showed (27) that the total human population on the entire territory of sub-Saharan Africa used in CassavaMap (7) increased by approximately 35% from 2014 to 2022; while the human population on a smaller territory located, for example, between 13° and 15° degrees of longitude and between 4° and 6° degrees of latitude (roughly the area considered in this paper), increased by approximately 17% from 2014 to 2022. Hence, rounding up, we assumed that the real area of cassava fields is +20% higher than recorded in CassavaMap.

To illustrate what the actual host landscape might be, we introduced additional scenarios into the original landscape, H, Fig. 2a, and created the following three alternative landscapes as surrogates for the actual landscape:

**Figure 2.**
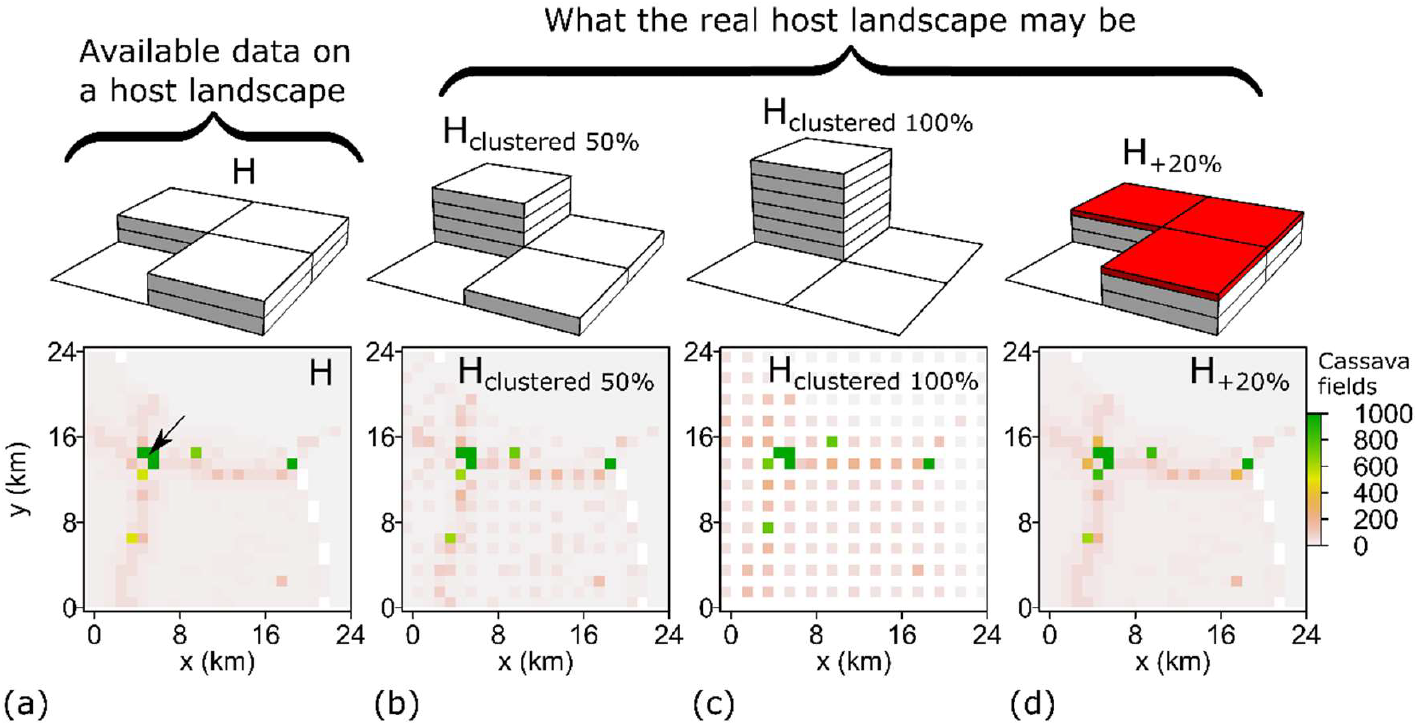
Host data and potential errors. **(a)** The landscape H represents the available data extracted from CassavaMap, Fig. 1. The small sample area shown here is the same area outlined by the blue boundary in Fig. 1. Black arrow in H denotes the location of a single initially infected cassava field (the same for all landscapes). Panels **(b), (c), (d)** illustrate different scenarios from H to account for potential errors in the mapped cassava landscape, illustrated on a 2 km by 2 km area shown above each landscape (also see Methods). Each tile represents a fixed number of fields, red tiles in (d) represent +20% in addition to two tiles underneath.

1. H_clustered 50%_ — To obtain the landscape denoted as ‘H_clustered 50%_’ we considered the original landscape H on a 2 km lattice, and in each 2 km by 2 km cell 50% of all cassava fields were first removed and then placed onto a North-Western 1 km by 1 km original raster cell. If the maximal capacity of that cell was reached, the excess fields were placed onto a North-Eastern cell, then South-Western and finally South-Eastern cells; see Fig. 2b. All fully occupied cells in the original unmodified landscape H were assumed to remain fully occupied.
2. H_clustered 100%_ — To obtain the landscape denoted as ‘H_clustered 100%_’ we followed the same procedure as for ‘H_clustered 50%_’ but using all cassava fields (i.e. 100% instead of 50%) in each 2 km by 2 km cell; see Fig. 2c.
3. H_+20%_ – We obtained the landscape denoted as ‘H_+20%_’ by adding 20% to the number of cassava fields present in each original 1 km by 1 km raster cell. If the maximum number of fields in any single 1 km by 1 km cell was reached because of adding new fields, then the excess of fields was distributed among eight nearest neighbouring cells by populating cells one-by-one up to the maximum level starting from the most occupied cells, see Fig. 2d.

### 2.3. The model of pathogen invasion and spread

The raster-based compartmental epidemiological SI (susceptible-infected) model of CBSV invasion and spread in sub-Saharan Africa was formulated in Godding *et al*. (5) and fitted to the data for CBSV spread in Uganda (5). The model (5) used a raster-based power-law dispersal kernel characterised by: the exponent α, the proportion *p* of dispersed inoculum that remains within the source (1 km by 1 km) cell, and the infection rate per contact density β. Here, using the approach of (18,28), we reformulated the raster-based model from Godding *et al*. (5) as an IBM in continuous space describing the spread of infection among discrete fields of cassava (treated as individual hosts within the IBM). An infected field infects susceptible fields at distance *x* with rate given by the product of the parameter β and a rotationally symmetric dispersal kernel, *b*(*x*). We applied the power-law dispersal kernel and selected the following parameter values from the posterior distribution from Godding *et al*. (5): α = 3.75, *p* = 0.12, and β = 10 × exp(6), see Fig. 3a; note, the parameter β in this paper is 10 times smaller than β in (5) because here it is measured in units [number of fields per area]^-1^× time^-1^ instead of units [number of raster cells per area]^-1^× time^-1^ used in (5). We considered CBSV invasion and spread over the four landscapes described above, i.e. H, H_clustered 50%_, H_clustered 100%_, and H_+20%_. Epidemics were simulated using the `ModelSimulator` software presented by Cornell *et al*. (29).

**Figure 3.**
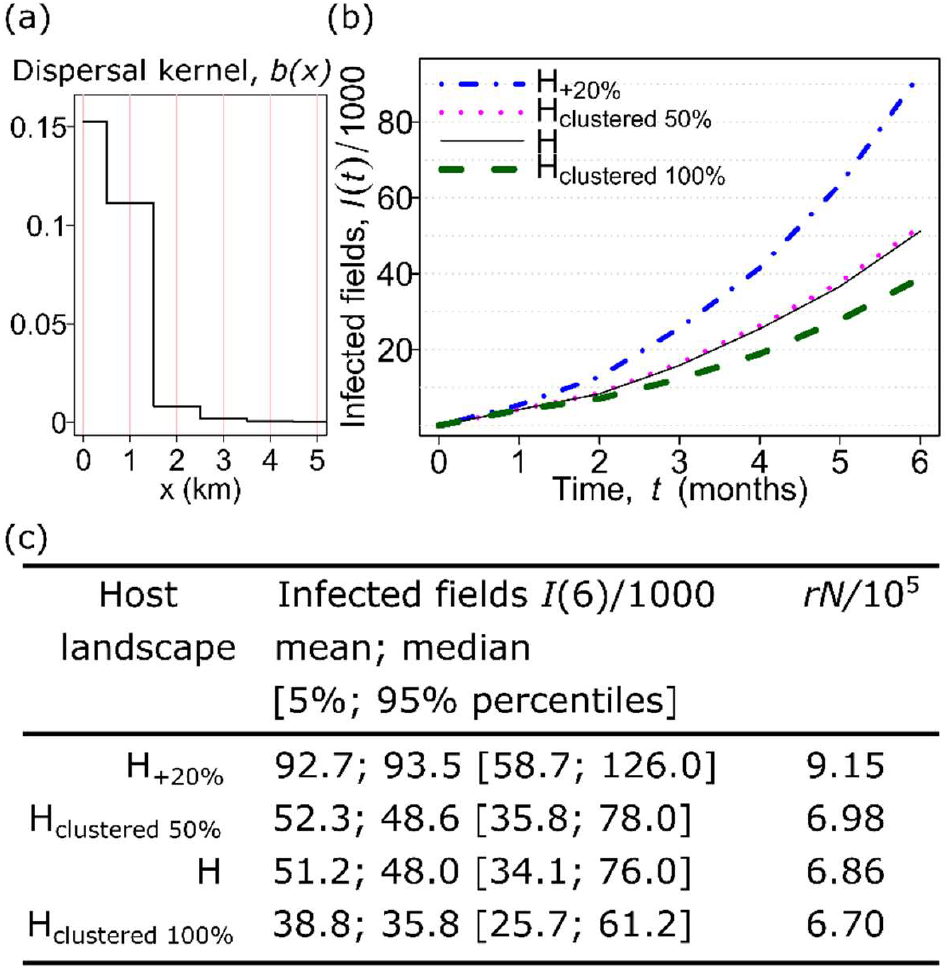
Impact of potential errors in host data on results of epidemic modelling. **(a)** Dispersal kernel *b*(*x*) of a pathogen (CBSV), sampled from results of parameter estimation by Godding *et al*. (5), see Methods. **(b)** Mean number *I*(*t*) of infected fields of cassava at *t* = 6 months after the start of an epidemic obtained from 1000 computer simulations for each landscape. **(c)** Mean, median and percentiles of results of computer simulations shown together with estimates of infection rate *r* from Equation (2.1) multiplied by the total number *N* of cassava fields estimated for corresponding 24 km by 24 km areas from Fig. 1.

### 2.4. Estimating the impact of potential inaccuracies in crop data on invasion of a pathogen

To determine the impact of potential inaccuracies in the cassava landscape on pathogen invasion using computer simulations, we considered the difference between the mean number of infected fields, *I*(*t*), at *t* = 6 months after the start of an epidemic in the alternative landscape, H_*Alt*_, and the (360 km by 360 km) landscape H derived from CassavaMap, i.e.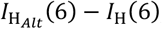, where the alternative landscape is one of the following: H_clustered 50%_, H_clustered 100%_, or H_+20%_ (Fig. 3b-c). For simplicity, in all cases, the same initial location of primary infection is used (Fig. 2a).

To determine the impact of potential inaccuracies on CBSV invasion analytically, we used an approximation (18) for the infection rate at which susceptible fields (i.e. individual hosts in an IBM) become infected at the start of an epidemic:

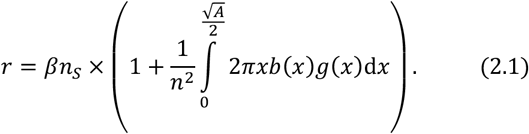

Here, *n*_S_ (or *n*) represents the spatial density of susceptible (or susceptible and infected) fields; β is the infection rate per contact density and *b*(*x*) is a pathogen dispersal kernel; *g*(*x*) is the spatial autocovariance of fields containing the host crop across the landscape of interest, *A*, and takes values on the interval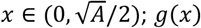 is also known as a second order spatial cumulant (e.g. see (29)). Equation (2.1) provides a spatially local estimate of the infection rate, *r*, on the local square area *A*. Denoting the total number of fields within the area *A* as *N*, we have *n* = *N*/*A*. We assumed that a pathogen was introduced by a single infected field randomly selected from all susceptible fields within area *A*, therefore *n*_S_ = (*N* − 1)/*A*. Note, that Equation (2.1) can be used for estimates of the basic reproduction number *R*_0_ in the corresponding SIR compartmental epidemiological model (e.g. see (18,30)): *R*_0_ can be approximated as the ratio of *r* and *μ*, i.e. *R*_0_ = *r*/*μ*, where *μ* is the removal or recovery rate of infected fields.

We compared the impact of alternative landscapes on epidemic invasion using the analytical approximation in Equation (2.1) and a metric, for the difference,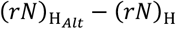, between *rN* in an alternative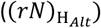and landscape H derived from CassavaMap ((*rN*)_H_). Positive (negative) values of this metric mean that an epidemic on the alternative (i.e. presumed real) landscape on average is larger (smaller) than an epidemic modelled on the landscape H.

As the quantity *rN* is measured locally, to select the size of the local area to estimate *rN* we used the results of Suprunenko *et al*. (18) where for the dispersal kernel *b*(*x*) (cf Fig. 3a) it was shown that the calculation of *r* on an area *A* = 8 × 8 km^2^ provides a better agreement with estimates from computer simulations than on a larger area, *A* = 24 × 24 km^2^. Therefore, we considered a host landscape on a square lattice with mesh size 8 km for which we estimated values of *n*_*S*_, *n, g*(*x*), *r* and *N* in each 8 km by 8 km cell.

### 2.5. Finding areas where inaccuracies in host data have the strongest impact on the modelled epidemic invasion

Here, we aim to construct a spatially resolved map of the degree of impact of potential errors in the host data on modelled epidemics. To achieve the aim using computer simulations would involve time-consuming efforts requiring a large number of computer simulations using different initial conditions and different implementations of potential errors. Instead, we adapt the analytical approach (based on Equation (2.1)) to identify spatial reconfigurations of a host landscape that provide the strongest deceleration of an invading pathogen (18). First, we assumed that H_+20%_ (Fig. 4a) is a more realistic surrogate of the actual landscape than H (i.e. it is a more accurate landscape than H), and that the effect of changing the host landscape could be shown on a square lattice with mesh size 2 km. Therefore, we divided each 8 km by 8 km cell (on the lattice used to calculate local *r*) into 2 km by 2 km cells.

**Figure 4.**
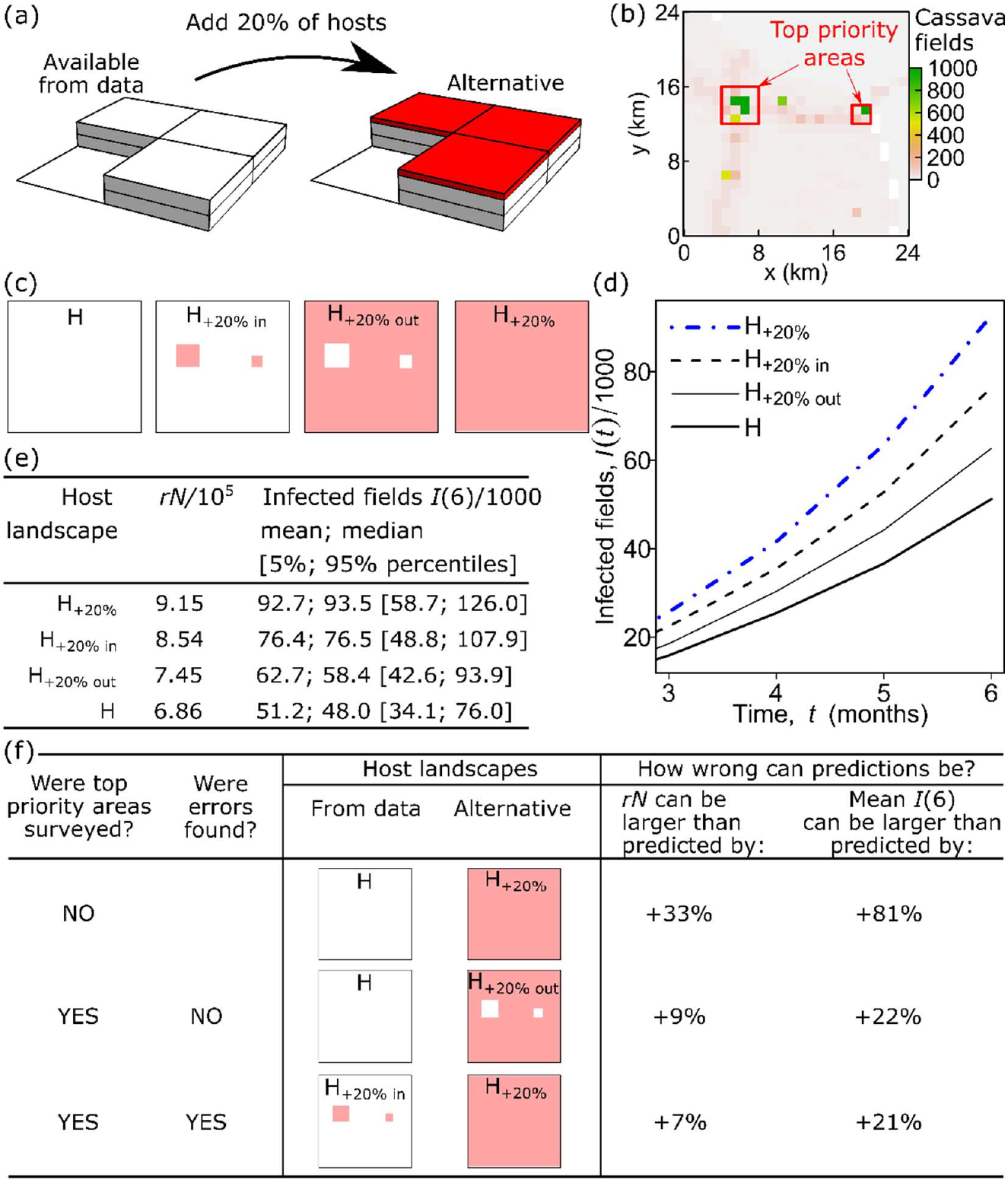
Finding areas where spatial host data should be refined to improve accuracy in crop disease modeling. **(a)** Presume that the alternative landscape, H_+20%_, is a realistic surrogate for the actual landscape (cf Fig. 1d), i.e. H_+20%_ has +20% more cassava fields than landscape H extracted from available data; see Methods for details. **(b)** Spatial prioritisation for data refinement: in each 24 km by 24 km cell within the entire landscape we identified five 2 km by 2 km cells with the highest impact on epidemic spread and denoted them as “top priority areas”, outlined by the red boundary. A small sample area shown here is the same area outlined by the blue boundary in Fig. 1. **(c)** The four landscapes in which: no fields are added (H); fields are added inside (H_+20% in_), or outside top priority areas (H_+20% out_) or across the domain (H_+20%_), subject to a maximum increase of 20% over the default map. **(d)** Mean number of infected fields obtained from 1000 computer simulations. **(e)** Numerical values for mean, median and percentiles of infected fields, together with estimates of quantities *rN* for each landscape. **(f)** The effect of spatial prioritisation and subsequent data refinement within identified top priority areas on the accuracy of epidemic model predictions, see Results for details. The computer code and data including entire landscapes and the map of top priority areas used in this work are available online from Figshare (24,27).

Calculating the product of the local infection rate *r* (calculated on 8 km by 8 km cells as described above) and the number *N*_2km_ of hosts in each 2 km-by-2 km cell on the H_+20%_ and H landscapes, we obtained the rasterized map of values 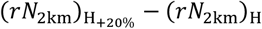. The areas with the largest values of the difference 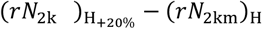 are areas where the selected inaccuracy has the strongest impact on an epidemic invasion.

### 2.6. Comparing the analytical solution with computer simulations

We used computer simulations to check the effect on epidemic dynamics from a change in the landscape due to spatial prioritisation provided by the analytical solution. We considered the situation when the landscape was divided into relatively small regions (analogous to local administrative regions) where spatial prioritisation was based on the data within that region independently from other local regions. Therefore, we considered the entire landscape on a square lattice with mesh size 24 km, within the overall domain of 336 km by 336 km, and within each 24 km by 24 km cell we used the algorithm presented above to select the top five 2 km by 2 km cells with the largest values of the difference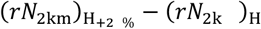. We refer to selected cells as “top priority areas” for data refinement (Fig. 4b shows top priority areas within a single 24 km by 24 km cell as an example; top priority areas identified simultaneously in all 24 km by 24 km cells within the entire landscape are available online from Figshare (24)). Next, we constructed two new landscapes in addition to landscapes H and H_+20%_: (i) the landscape denoted “H_+20%, in_” was obtained from H by introducing +20% of hosts *only inside* the selected five 2 km by 2 km cells *in each* 24 km by 24 km area; (ii) the landscape denoted “H_+20%, out_” was obtained from H by introducing +20% of hosts in all cells *only outside* the selected five cells *in each* 24 km by 24 km area; the difference between the four landscapes on a sample 24 km by 24 km area is illustrated in Fig. 4c. If the area with the highest impact is identified correctly, then it is expected that the epidemic on landscape H_+20%, in_ should be closer (in terms of the mean number of infected fields) to the epidemic on landscape H_+20%_ than on landscape H; similarly, the epidemic on H_+20%, out_ should be closer to the epidemic on H than on H_+20%_. Using computer simulations of the IBM together with analytical estimates describe above, we compared the characteristics of an epidemic invasion on the four landscapes described in this section, Fig. 4d-e.

## 3. Results

Computer simulations of the IBM of a potential CBSV epidemic using the selected dispersal kernel (Fig. 3a) demonstrated that potential inaccuracies in the host data (Fig. 2) can influence epidemic invasion strongly. In particular, we found that the mean number of infected cassava fields could be 24% less on a more clustered landscape compared with the H landscape calculated from CassavaMap (i.e. -24% in H_clustered 100%_ relative to H, Fig. 3b-c). By contrast up to 81% more fields were infected compared with H when allowance was made for up to 20% missing fields on CassavaMap (i.e. +81% in H_+20%_ relative to H, Fig. 3b-c). Recall that the alternative landscapes are designed to allow for inaccuracies in H: H_clustered 100%_ allows for failure to account for clustering if fields were over-dispersed amongst CassavaMap cells from coarse production data; the H_+20%_ landscape allows for growth in human population density in the decade since the raw data for CassavaMap were collated.

We used the product, *rN*, of the local infection rate multiplied by the local number of hosts as an analytical approximation to map the impact from inaccuracies in the host data on the modelled epidemic dynamics. Geographic areas with the largest impact were indicated as priorities for data refinement efforts (i.e. where additional data on the spatial distribution of fields of susceptible crops would be valuable).

To verify the analytical method of spatial prioritisation, we tested the effect of data refinement in top priority areas. To compare the impact of inaccuracies in the identified top priority areas and in the rest of the landscape, we constructed two additional landscapes where potential inaccuracies were adjusted for by changing the number of host fields either only inside or only outside those top priority areas (Fig. 4c). Calculated values of *rN* and computer simulations of CBSV spread confirmed that the impact of inaccuracies in the selected top priority area was indeed stronger than the impact from the rest of the landscape (Fig. 4d-e).

Finally, we estimated how the accuracy of epidemic model predictions would change because of spatial prioritisation and subsequent data refinement within the top priority areas. Using the example studied in Fig. 4a-e, we focused on the following three cases, Fig. 4f:

i. in the first case, we assumed that top priority areas were not identified and therefore no additional data were collected. Therefore, H_+20%_ was considered as the most realistic surrogate for the actual landscape. In that case, the potential inaccuracy of the model predictions was +81% in the alternative landscape H_+20%_ as compared with the landscape H extracted from the data (CassavaMap), i.e. 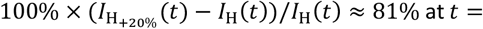6 months, where *I*(*t*) is the mean number of infected fields as inferred from computer simulations of the IBM.
ii. In the second case, we assumed that the top priority areas were identified, and the subsequent survey of the top priority areas showed no errors in the landscape H in those areas. Therefore, H_+20% out_ was considered as the most realistic surrogate for the actual landscape, i.e. inaccuracies could be present only outside the identified areas. In that case, the mean number of infected fields, *I*(6), was +22% larger than model predictions based on the landscape H.
iii. In the third case, we assumed that survey of the identified top priority areas confirmed the presumed inaccuracies, i.e. the number of hosts in those areas was larger by the presumed +20% than in the landscape H. In that case, the refined data described the landscape H_+20% in_ (i.e. inaccuracies were present only in the identified areas), and H_+20%_ was considered as the most realistic surrogate for the actual landscape. As a result, an epidemic on H_+20%_ was larger by +21% than predictions based on the refined data (H_+20% in_).

Note, in both cases (ii) and (iii) the identification and subsequent survey of the top priority areas reduced the potential deviation in model predictions for the variable *I*(6) by approximately four times (from 81% down to 22% or 21%, Fig.3f). A qualitatively similar reduction was also observed in the analytical estimate of *rN*: there was approximately four-fold reduction of deviations (from 33% down to 9%) when no errors were found in the data and approximately five-fold reduction of deviations (from 33% down to 7%) when the presumed errors in the data were confirmed. Thus, these findings suggest that the method based on the estimate of the quantity *rN* could be used to identify top priority areas for additional survey that would improve accuracy in epidemic modelling.

## 4. Discussion

We have presented a solution to the problem of identifying local geographic priorities for host data refinement efforts that would improve the accuracy of the number of infected fields of crops predicted by an epidemic spread model during the early stage of an epidemic (Fig. 4). As an example of an epidemic of a crop disease, we considered the spread of CBSV through an arbitrarily selected 336 km by 336 km cassava landscape (derived from CassavaMap (7)) in sub-Saharan Africa where the potential errors in host data (Fig. 2) can have a substantial effect on predictions of the number of infected fields (Fig. 3).

The use of an analytical approach enabled the identification of priority areas for host refinement in relation to potential impacts on epidemic spread. It would be practically unfeasible to derive an equivalent solution by computer simulation due to a large number of alternative configurations that would have to be considered: for example, in addition to multiple simulations for each initial condition in each landscape, one would need to consider many alternative landscapes where the selected type of inaccuracy is placed in all possible locations. Within the analytical approach we used the quantity *rN* because it has a number of characteristics that are important for the solution derived in this paper. The variable, *rN* is measured locally, i.e. within a selected local area; *rN* captures changes in both *r* (that is influenced by the pathogen dispersal kernel and the local spatial structure of the host landscape) and in *N*, the number of hosts in the local area; *rN* is an additive characteristic in the sense that *rN* characterizing a larger area is simply a sum of *rN* estimated from smaller areas within the larger one: therefore *rN* allows a direct comparison of contributions from different local areas. An additional advantage of the analytical approach over computer simulations is that it can be used to derive general insights into the dependence of the spatial prioritisation for data refinement on the interplay of several factors. These include different types of potential inaccuracies in host data as well as the spatial pattern, especially intercrop clustering, relative to the dispersal kernel of an invading pathogen.

For practical applications, it is important to stress that we assumed that all host crops within a landscape have the same probability of becoming infected at the start of the epidemic. Based on this assumption, areas with the strongest impact from the selected inaccuracy were identified as the top priority areas for data refinement. However, where there are local differences in the probability of crops becoming infected in certain areas, spatial prioritisation for data refinement would need to account for that heterogeneity. Differences could arise due to local use of pesticide, cultivation of partially resistant varieties or where environmental conditions differ. Extension of the analytical method to deal with this type of heterogeneities could be considered in future work.

In addition to the analytical approximation, Equation (2.1), used in this paper, there are other equivalent approximations for the infection rate as well as the closely related quantity, the basic reproduction number *R*_0_. Other approximations that are applicable to non-random host distribution (and therefore relevant to this work) have been derived, for example, in Bolker (31), North and Godfray (32), and more recently by van den Bosch *et al*. (33). As shown in Suprunenko *et al*. (18), the estimation of the infection rate according to Equation (2.1) differs from earlier estimates (31–33) only when calculated in a local area of a host landscape with low density of hosts, i.e. when the difference between the density of susceptible hosts, *n*_S_, and the total density, *n*, matters. In the case of cassava in sub-Saharan Africa, rural areas are often characterised (in CassavaMap) by a low number of cassava fields per unit area, therefore the selected approximation (Equation (2.1)) has a slightly higher accuracy as compared with alternative approximations (31–33). Other approximations for infection rate and the basic reproduction number, *R*_0_, would be useful in some specific cases of spatial distribution of hosts. For example, van den Bosch *et al*. (33) derived analytical expressions for *R*_0_ in regular host distributions generated by a Strauss process, and in spatially clustered distributions generated by a Neyman-Scott process. For random spatial host distributions, Suprunenko *et al*. (30) derived a spatially local approximation for *r*, and Wadkin *et al*. (34) improved the accuracy of the approximation for *r* and *R*_0_ by accounting for host depletion. Mikaberidze *et al*. (35) considered a single rectangular crop field for which they derived *R*_0_ from a system of integro-differential equations. In addition, *R*_0_ can be calculated by using all pairwise probabilities of a host infecting any other hosts, for example see Tildesley and Keeling (16) and references therein. Further work is needed to review the merits of the different approaches, ideally for real systems The challenge associated with spatial prioritisation for data collection or control and management efforts has attracted attention in recent studies of disease spread in agricultural systems. For example, when studying livestock disease with limited host data, Dawson *et al*. (10) modelled disease spread on the UK cattle trade network and showed that nodes with the highest number of livestock movements should be prioritised for data collection to get more accurate model predictions of an epidemic. Spatial prioritisation for data collection aiming to improve accuracy of spatial modelling of crop diseases is a relatively new topic that supplements a larger field of research on spatial prioritisation for surveillance for plant pests and pathogens (36–38) as well as spatial prioritisation of management efforts for greater crop yield (39).

Future work could develop the analytical approach presented here to address some open problems. While our analyses have assumed rotationally symmetrical dispersal kernels, crop pathogens often disperse anisotropically. Wheat stem rust (3) for example is dispersed anisotropically by wind, therefore, incorporating anisotropical dispersal of pathogens into methods of this paper could potentially help in refining wheat field data at various scales (4) to improve accuracy in epidemic modelling. As another example, the lack of spatial information within raster cells in rasterized host data with coarse spatial resolution can potentially be a source of large inaccuracy in raster-based epidemic models. Tildesley *et al*. (8) addressed this problem in optimal control of foot and mouth disease of cattle and showed that data on exact farm locations were not required and that using randomized aggregate county-scale data was sufficient when the model parameters could be re-fitted to the outbreak data on randomized locations. In addition, it was shown that missing spatial information within raster cells in rasterized host data can also be imputed by using land-cover maps (9) or can be predicted by using computational methods such as the Farm Location and Agricultural Production Simulator (11). However, investigation of the problem of the coarse spatial resolution in rasterized host data in a broader range of applied research questions would be valuable.

## Acknowledgements

The authors are grateful to Dr Alison Scott-Brown, Dr Renata Retkute and Dr Ruairi Donnelly for helpful comments on the manuscript.

## References

1. Cunniffe NJ, Koskella B, E. Metcalf CJ, Parnell S, Gottwald TR, Gilligan CA. Thirteen challenges in modelling plant diseases. Epidemics [Internet]. 2015 Mar [cited 2024 Jun 26];10:6–10. Available from: https://linkinghub.elsevier.com/retrieve/pii/S1755436514000309

2. Meyer M, Bacha N, Tesfaye T, Alemayehu Y, Abera E, Hundie B, et al. Wheat rust epidemics damage Ethiopian wheat production: A decade of field disease surveillance reveals national-scale trends in past outbreaks. Zhang A, editor. PLoS ONE [Internet]. 2021 Feb 3 [cited 2024 Aug 8];16(2):e0245697. Available from: https://dx.plos.org/10.1371/journal.pone.0245697

3. Bradshaw CD, Thurston W, Hodson D, Mona T, Smith JW, Millington SC, et al. Irrigation can create new green bridges that promote rapid intercontinental spread of the wheat stem rust pathogen. Environ Res Lett [Internet]. 2022 Nov 1 [cited 2024 Jul 7];17(11):114025. Available from: https://iopscience.iop.org/article/10.1088/1748-9326/ac9ac7

4. Blasch G, Alemayehu Y, Lesne L, Wolter J, Taymans M, Tesfaye T, et al. Ethiopian Crop Type 2020 (EthCT2020) dataset: Crop type data for environmental and agricultural remote sensing applications in complex Ethiopian smallholder wheat-based farming systems (Meher season 2020/21). Data in Brief [Internet]. 2024 Jun [cited 2024 Jul 7];54:110427. Available from: https://linkinghub.elsevier.com/retrieve/pii/S2352340924003962

5. Godding D, Stutt Rojh, Alicai T, Abidrabo P, Okao-Okuja G, Gilligan CA. Developing a predictive model for an emerging epidemic on cassava in sub-Saharan Africa. Sci Rep [Internet]. 2023 Aug 3 [cited 2024 Mar 11];13(1):12603. Available from: https://www.nature.com/articles/s41598-023-38819-x

6. Hassall KL, Alonso Chávez V, Sint H, Helps JC, Abidrabo P, Okao-Okuja G, et al. Validating a cassava production spatial disaggregation model in sub-Saharan Africa. Alleyne AT, editor. PLoS ONE [Internet]. 2024 Nov 5 [cited 2024 Dec 2];19(11):e0312734. Available from: https://dx.plos.org/10.1371/journal.pone.0312734

7. Szyniszewska AM. CassavaMap, a fine-resolution disaggregation of cassava production and harvested area in Africa in 2014. Sci Data [Internet]. 2020 May 27 [cited 2024 Mar 11];7(1):159. Available from: https://www.nature.com/articles/s41597-020-0501-z

8. Tildesley MJ, House TA, Bruhn MC, Curry RJ, O’Neil M, Allpress JLE, et al. Impact of spatial clustering on disease transmission and optimal control. Proc Natl Acad Sci USA [Internet]. 2010 Jan 19 [cited 2024 Jul 1];107(3):1041–6. Available from: https://pnas.org/doi/full/10.1073/pnas.0909047107

9. Tildesley MJ, Ryan SJ. Disease Prevention versus Data Privacy: Using Landcover Maps to Inform Spatial Epidemic Models. Salathé M, editor. PLoS Comput Biol [Internet]. 2012 Nov 1 [cited 2024 Oct 2];8(11):e1002723. Available from: https://dx.plos.org/10.1371/journal.pcbi.1002723

10. Dawson PM, Werkman M, Brooks-Pollock E, Tildesley MJ. Epidemic predictions in an imperfect world: modelling disease spread with partial data. Proc R Soc B [Internet]. 2015 Jun 7 [cited 2024 Jul 1];282(1808):20150205. Available from: https://royalsocietypublishing.org/doi/10.1098/rspb.2015.0205

11. Burdett CL, Kraus BR, Garza SJ, Miller RS, Bjork KE. Simulating the Distribution of Individual Livestock Farms and Their Populations in the United States: An Example Using Domestic Swine (Sus scrofa domesticus) Farms. Van Boven M, editor. PLoS ONE [Internet]. 2015 Nov 16 [cited 2024 Oct 2];10(11):e0140338. Available from: https://dx.plos.org/10.1371/journal.pone.0140338

12. Sellman S, Tildesley MJ, Burdett CL, Miller RS, Hallman C, Webb CT, et al. Realistic assumptions about spatial locations and clustering of premises matter for models of foot-and-mouth disease spread in the United States. Kao R, editor. PLoS Comput Biol [Internet]. 2020 Feb 20 [cited 2024 Oct 2];16(2):e1007641. Available from: https://dx.plos.org/10.1371/journal.pcbi.1007641

13. Vizzari M, Lesti G, Acharki S. Crop classification in Google Earth Engine: leveraging Sentinel-1, Sentinel-2, European CAP data, and object-based machine-learning approaches. Geo-spatial Information Science [Internet]. 2024 Apr 25 [cited 2024 Jul 1];1–16. Available from: https://www.tandfonline.com/doi/full/10.1080/10095020.2024.2341748

14. Alvarez-Vanhard E, Corpetti T, Houet T. UAV & satellite synergies for optical remote sensing applications: A literature review. Science of Remote Sensing [Internet]. 2021 [cited 2024 Jul 9];3:100019. Available from: https://linkinghub.elsevier.com/retrieve/pii/S2666017221000067

15. Kaivosoja J, Hautsalo J, Heikkinen J, Hiltunen L, Ruuttunen P, Näsi R, et al. Reference Measurements in Developing UAV Systems for Detecting Pests, Weeds, and Diseases. Remote Sensing [Internet]. 2021 Mar 24 [cited 2024 Jul 9];13(7):1238. Available from: https://www.mdpi.com/2072-4292/13/7/1238

16. Tildesley MJ, Keeling MJ. Is R0 a good predictor of final epidemic size: Foot-and-mouth disease in the UK. Journal of Theoretical Biology [Internet]. 2009 Jun [cited 2024 Oct 2];258(4):623–9. Available from: https://linkinghub.elsevier.com/retrieve/pii/S0022519309000824

17. Delamater PL, Street EJ, Leslie TF, Yang YT, Jacobsen KH. Complexity of the Basic Reproduction Number (R 0). Emerg Infect Dis [Internet]. 2019 Jan [cited 2024 Oct 6];25(1):1–4. Available from: http://wwwnc.cdc.gov/eid/article/25/1/17-1901_article.htm

18. Suprunenko YF, Cornell SJ, Gilligan CA. Predicting the effect of landscape structure on epidemic invasion using an analytical estimate for infection rate. R Soc Open Sci [Internet]. 2024;11:240763. Available from: 10.1098/rsos.240763

19. Nweke FI, Lynam JK, Spencer DSC. The Cassava Transformation: Africa’s Best-Kept Secret [Internet]. Michigan State University Press; 2002. 273 p. Available from: https://www.jstor.org/stable/10.14321/j.ctt7ztc0t

20. Patil BL, Legg JP, Kanju E, Fauquet CM. Cassava brown streak disease: a threat to food security in Africa. Journal of General Virology [Internet]. 2015;96:956–68. Available from: https://pubmed.ncbi.nlm.nih.gov/26015320/

21. Legg JP, Jeremiah SC, Obiero HM, Maruthi MN, Ndyetabula I, Okao-Okuja G, et al. Comparing the regional epidemiology of the cassava mosaic and cassava brown streak virus pandemics in Africa. Virus Research [Internet]. 2011 Aug [cited 2024 Jun 28];159(2):161–70. Available from: https://linkinghub.elsevier.com/retrieve/pii/S0168170211001547

22. Alicai T, Omongo CA, Maruthi MN, Hillocks RJ, Baguma Y, Kawuki R, et al. Re-emergence of cassava brown streak disease in Uganda. Plant Disease [Internet]. 2007 Jan [cited 2024 Mar 12];91(1):24–9. Available from: https://apsjournals.apsnet.org/doi/10.1094/PD-91-0024

23. Muhindo H, Wembonyama F, Yengele O, Songbo M, Tata-Hangy W, Sikirou M, et al. Optimum time for harvesting cassava tubers to reduce losses due to cassava brown streak disease in northeastern DRC. JAS [Internet]. 2020 Apr 15 [cited 2024 Mar 11];12(5):70. Available from: http://www.ccsenet.org/journal/index.php/jas/article/view/0/42480

24. Suprunenko YF, Gilligan CA. Data from: ‘Where to refine spatial data to improve accuracy in crop disease modelling: an analytical approach for mapping the impact of potential errors in crop data with examples for cassava’. 2024. (In review).

25. Bright E, Rose A, Urban M. LandScan Global 2014 [Data set] [Internet]. Oak Ridge National Laboratory; 2015. Available from: 10.48690/1524209

26. Sims K, Reith A, Bright E, Kaufman J, Pyle J, Epting J, et al. LandScan Global 2022 [Data set] [Internet]. Oak Ridge National Laboratory; 2023. Available from: 10.48690/1529167

27. Suprunenko YF, Gilligan CA. Computer code for: ‘Where to refine spatial data to improve accuracy in crop disease modelling: an analytical approach for mapping the impact of potential errors in crop data with examples for cassava’. 2024. (In review).

28. Suprunenko YF, Cornell SJ, Gilligan CA. Computer code for: Predicting the effect of landscape structure on epidemic invasion using an analytical estimate for infection rate [Internet]. Figshare; 2024. Available from: 10.6084/m9.figshare.25804810

29. Cornell SJ, Suprunenko YF, Finkelshtein D, Somervuo P, Ovaskainen O. A unified framework for analysis of individual-based models in ecology and beyond. Nat Commun [Internet]. 2019 Oct 17 [cited 2024 Mar 11];10(1):4716. Available from: https://www.nature.com/articles/s41467-019-12172-y

30. Suprunenko YF, Cornell SJ, Gilligan CA. Analytical approximation for invasion and endemic thresholds, and the optimal control of epidemics in spatially explicit individual-based models. J R Soc Interface [Internet]. 2021 Mar [cited 2024 Mar 11];18(176):rsif.2020.0966, 20200966. Available from: https://royalsocietypublishing.org/doi/10.1098/rsif.2020.0966

31. Bolker B. Analytic models for the patchy spread of plant disease. Bulletin of Mathematical Biology [Internet]. 1999 Sep [cited 2024 Mar 11];61(5):849–74. Available from: http://link.springer.com/10.1006/bulm.1999.0115

32. North AR, Godfray HCJ. The dynamics of disease in a metapopulation: The role of dispersal range. Journal of Theoretical Biology [Internet]. 2017 Apr [cited 2024 Mar 11];418:57–65. Available from: https://linkinghub.elsevier.com/retrieve/pii/S0022519317300371

33. van den Bosch F, Helps J, Cunniffe NJ. The basic-reproduction number of infectious diseases in spatially structured host populations. Oikos [Internet]. 2024 Jun 18 [cited 2024 Jul 7];e10616. Available from: 10.1111/oik.10616

34. Wadkin LE, Holden J, Ettelaie R, Holmes MJ, Smith J, Golightly A, et al. Estimating the reproduction number, R 0, from individual-based models of tree disease spread. Ecological Modelling [Internet]. 2024 Mar [cited 2024 Mar 11];489:110630. Available from: 10.1016/j.ecolmodel.2024.110630

35. Mikaberidze A, Mundt CC, Bonhoeffer S. Invasiveness of plant pathogens depends on the spatial scale of host distribution. Ecological Applications [Internet]. 2016 Jun [cited 2024 Mar 11];26(4):1238–48. Available from: https://esajournals.onlinelibrary.wiley.com/doi/10.1890/15-0807

36. Tuomola J, Yemshanov D, Huitu H, Hannunen S. Mapping risks of pest invasions based on the spatio-temporal distribution of hosts. MBI [Internet]. 2018 [cited 2024 Oct 2];9(2):115–26. Available from: http://www.reabic.net/journals/mbi/2018/Issue2.aspx

37. Mastin AJ, Gottwald TR, Van Den Bosch F, Cunniffe NJ, Parnell S. Optimising risk-based surveillance for early detection of invasive plant pathogens. Perrings C, editor. PLoS Biol [Internet]. 2020 Oct 12 [cited 2024 Nov 13];18(10):e3000863. Available from: https://dx.plos.org/10.1371/journal.pbio.3000863

38. Soubeyrand S, Estoup A, Cruaud A, Malembic-Maher S, Meynard C, Ravigné V, et al. Building integrated plant health surveillance: a proactive research agenda for anticipating and mitigating disease and pest emergence. CABI Agric Biosci [Internet]. 2024 Aug 17 [cited 2024 Dec 2];5(1):72. Available from: https://cabiagbio.biomedcentral.com/articles/10.1186/s43170-024-00273-8

39. Buddenhagen CE, Xing Y, Andrade-Piedra JL, Forbes GA, Kromann P, Navarrete I, et al. Where to Invest Project Efforts for Greater Benefit: A Framework for Management Performance Mapping with Examples for Potato Seed Health. Phytopathology® [Internet]. 2022 Jul [cited 2024 Jul 3];112(7):1431–43. Available from: https://apsjournals.apsnet.org/doi/10.1094/PHYTO-05-20-0202-R

